# HIV-1 VPR TARGETS BCOR FOR DEGRADATION TO COUNTER POLYCOMB-DEPENDENT PROVIRAL SILENCING

**DOI:** 10.64898/2026.11.26.769605

**Authors:** Maria Carmen Romero-Medina, Isa Muñoz-Arias, Jérémy A. Ferreira Barbosa, Tram N.Q. Pham, Éric A. Cohen

## Abstract

Chromatin remodeling complexes such as the Polycomb Repressive Complex (PRC) are recruited to the HIV-1 promoter during latency. Using BioID, we previously identified multiple PRC subunits as potential HIV-1 Vpr-interacting partners. Here, we show that HIV-1 Vpr, but not its paralog Vpx encoded by SIVsmm/HIV-2, targets the non-canonical PRC1.1 core component, BCL6 co-repressor (BCOR), for proteasomal degradation. Vpr-dependent depletion of BCOR is observed during infection of T-cells and can be induced by virion-associated Vpr prior to early step leading to viral DNA integration and provirus expression. BCOR degradation is independent from the G2/M-arrest activity induced by Vpr but requires the recruitment of the CRL4^DCAF1^ E3 ubiquitin ligase by Vpr. Using T-cell models of HIV latency (Jurkat E4 and 2D10), we further show that BCOR is recruited to the LTR of latent proviruses along with the polycomb group RING finger protein 1 (PCGF1) and H2AK119ub prototypical silencing marks are detectable in this context. ShRNA-directed BCOR knockdown reactivates HIV-1 expression in 2D10 cells, and this reactivation is phenocopied upon Vpr expression. Overall, our results identify PRC1.1 as the non-canonical complex involved in PRC-dependent silencing of the HIV-1 LTR and further reveal that Vpr counteracts HIV-1 transcriptional silencing through BCOR degradation likely to achieve optimal viral expression and propagation during infection.

**IMPORTANCE:** Antiretroviral therapy effectively suppresses HIV-1 replication but does not affect the persistent latent reservoir, which remains the primary obstacle to a cure. Understanding the HIV-host interactions that govern viral latency and persistence is therefore essential for designing strategies that reduce the size and activity of this reservoir. Here we report that Vpr, an accessory protein encoded by HIV-1, counteracts polycomb repressive complex (PRC)-dependent silencing of integrated provirus expression by depleting BCOR, a core component of the PRC1 subunit, which is responsible for ubiquitination of histones. These findings provide insights into how HIV-1 Vpr modulates a key host epigenetic pathway involved in HIV-1 latency and shed to light a host factor targeted by HIV that could be exploited to design approaches aimed at making HIV-1 reservoirs more visible for clearance by the host immune system.

## INTRODUCTION

Integration of the reverse transcribed viral genome into host chromatin is a defining feature of retroviruses that is central to their replication and ability to establish persistent infections. While human immunodeficiency virus type 1 (HIV-1) infection of activated CD4^+^ T cells typically results in high-level expression of the integrated proviral DNA, it is well recognized that the chromatin configuration at the site of provirus integration governs the extent of transcriptional activation of the integrated provirus (1–4). Indeed, cellular chromatin modifiers and remodelers, which post-translationally control the modification of histones by acetylation, methylation, and ubiquitination, contribute to chromatin compaction and viral silencing, notably in dense and transcriptionally silenced chromatin regions such as heterochromatin (5). While host factors that silence proviral transcription in CD4^+^ memory T cells help infected cells escape elimination by the host immune system and antiviral drugs (6), these same factors must, however, be counteracted for optimal HIV propagation during infection.

Previous studies have identified a role for the human silencing hub (HUSH) complex in the heterochromatinization and silencing of newly integrated lentiviruses, a process that was shown to be counteracted by the Vpx and Vpr accessory proteins encoded by diverse primate immunodeficiency viruses with the notable exception of HIV-1 Vpr (7–9). More recently, the structural maintenance of chromosome 5/6 complex localization factor 2 (SLF2) was found to specifically repress extrachromosomal unintegrated HIV-1 provirus expression by DNA-based compaction silencing, a restriction that was also antagonized by Vpr from divergent primate lentiviral lineages, including HIV-1 and HIV-2 (10, 11). Therefore, it seems that primate immunodeficiency viruses have evolved to evade host silencing mechanisms that distinctly act on the expression of unintegrated and integrated proviral DNA through their accessory proteins.

HIV-1 Vpr is a virion-incorporated accessory protein that is conserved across all HIV and simian immunodeficiency virus (SIV) lineages (12). Previous studies have shown that Vpr induces a G2/M cell cycle arrest during infection of dividing cells (13, 14). Study of the mechanism underlying this process uncovered that Vpr recruits the host Cul4-DDB1 cullin-Ring E3 ubiquitin ligase complex via the adaptor protein DCAF1 (CRL4^DCAF1^) to induce the proteasome-mediated degradation of target proteins (13, 15, 16). Among these host targets are proteins involved in post DNA-replication repair, such as uracil-DNA glycosylase (UNG2) (17), the double strand break repair and Holliday junction resolvase MUS81-EME1 (18), the helicase-like transcription factor, HLTF, (19, 20) and Exo1, a nuclease involved in double strand break DNA repair (21). However, unbiased proteomic analyses revealed that HIV-1 Vpr induces a global remodeling of the cellular proteome and highlighted potential interactions with a wide variety of biological pathways that could lead to G2/M cell cycle arrest, with several also having the potential to alter chromatin metabolism and silence proviral DNA (22–24). Indeed, in addition to SLF2 and the HUSH complex, Vpr was reported to target subunits from various chromatin remodeling/modifying complexes for degradation, including the transcriptional repressors and adaptors, CTIP2, zinc finger and G-patch domain-containing (ZIP) and sZIP of the NuRD chromatin remodeling complex, the methylcytosine dioxygenase TET2 and the histone deacetylase SIRT7, thereby implicating Vpr as a potential epigenetic regulator of host and proviral gene expression (25, 26) (27, 28).

In this context, we previously conducted a Vpr proximity-dependent biotin identification (BioID) analysis that identified subunits of the Polycomb Repressive Complex (PRC) as potential Vpr-interacting partners (24). The PRC system is associated with transcriptional silencing of heterochromatin and consists of two distinct groups of complexes with different enzymatic activities; PRC1, which contains the histone ubiquitin ligases, really interesting gene 1A or B (RING1A/B) proteins, and PRC2 complexes, which includes the methyltransferases enhancer of zeste homolog 1 or 2 (EZH1/2) (29) (30) (31) (32).

It has become clear that various mutually exclusive PRC1 and PRC2 variants exist with PRC1 assembling canonical (c) and non-canonical (nc) complexes depending on whether they contain a chromodomain box protein (CBX) or one of the two homologous proteins, RING1 and YY1 binding proteins (RYBP) and YY1-associated factor (YAF2), respectively (33, 34). A unique feature of cPRC1 variants is their dependency on PRC2-deposited H3K27me^3^ for chromatin occupancy, a property not shared by ncPRC1 (29, 33, 35). PRC1 complexes are also defined according to which of the six Polycomb group ring finger (PCGF1-6) proteins they contain, with PCGF2 and PCGF4 being able to form ncPRC1 or cPRC1, and the rest of the PCGF proteins only assembling ncPRC1 complexes (ncPRC1.1-1.6 vs cPRC1.2 and cPRC1.4) (29, 33, 35).

Increasing evidence suggests a critical role for PRC in the silencing of integrated HIV-1 provirus in latently infected CD4^+^ T cells. Using several T cell models of HIV latency, multiple PRC2.2 subunits, including EZH2, have been shown to occupy and repress the HIV-1 LTR (36) (37–39). The core component of PRC1.4, PCGF4 (also named BMI1), is also necessary for the maintenance and initiation of HIV-1 latency (37, 40, 41). For instance, PCGF4 occupancy of the LTR has been demonstrated to be dependent on EZH2 expression suggesting that PCGF4 is indeed forming a canonical complex (41). Additionally, the involvement of the transcription factor Ying Yang 1 (YY1), a signature component of ncPRC1 complexes, in inhibiting HIV-1 LTR-directed expression, strongly suggests that an unidentified ncPRC1 complex also contributes to PRC-dependent HIV-1 silencing (37, 42, 43). Interestingly, among the PRC subunits detected in close proximity to Vpr in our previously reported BioID analysis (24) was the ubiquitously expressed B-cell lymphoma 6 (BCL6) protein co-repressor (BCOR), a core component of the ncPRC1.1 complex (35, 44).

In this study, we investigated the role for HIV-1 Vpr in counteracting PRC-dependent silencing of integrated proviral DNA expression via targeting of the ncPRC1.1 core component BCOR. We show that HIV-1 Vpr specifically targets BCOR for degradation via a process that requires the recruitment of the CRL4^DCAF1^ ubiquitin ligase and an active proteasome. Using ChIP-qPCR analysis, we further show that BCOR and PCGF1, another core component of ncPRC1.1, and accompanying PRC-directed histone silencing marks occupy the HIV-1 LTR of latent integrated proviral DNA in T-cell models of latency (Jurkat E4 and 2D10). Downmodulation of BCOR by shRNA reactivates integrated HIV-1 DNA expression in these models of latency and this reactivation is also observed upon transduction of Vpr using lentiviral vectors. By degrading BCOR, HIV-1 Vpr counteracts the PRC-dependent silencing of integrated proviral DNA, allowing optimal viral expression and propagation during infection.

## RESULTS

### HIV-1 Vpr but not HIV-2/SIV_smm_ Vpx expression promotes BCOR depletion

Our previous work implicated BCOR amongst other PRC subunits as potential interacting partners of HIV-1 Vpr (24). Thus, to test whether Vpr depletes any of the PRC subunits identified in the BioID analysis, or proteins known to recruit BCOR to promoters, namely BCL6 and the demethylase KDM2B (29, 35), we used a HeLa Flp-In T-Rex system, which expresses hemagglutinin (HA)-tagged HIV-1_Bru_ Vpr upon induction with doxycycline (24). First, we show that HLTF, a well-established target of Vpr, was depleted only when Vpr was induced (Figure 1A; left panel) (20, 45), confirming that Vpr is functional. Over the course of 24 h of doxycycline-induced Vpr expression, BCOR levels were consistently reduced in comparison to uninduced cells. BCOR was the only PRC-related endogenous protein detected in the BioID analysis that was depleted by Vpr; PRC components such as RING1B, PCGF6 and SCML1 did not show any expression variation in the presence of Vpr (Figure 1 A; both panels). Furthermore, neither EZH2, a PRC2 component involved in HIV latency, nor BCL6 or KDM2B, were affected by Vpr expression (Figure 1A; both panels).

**Figure 1.**
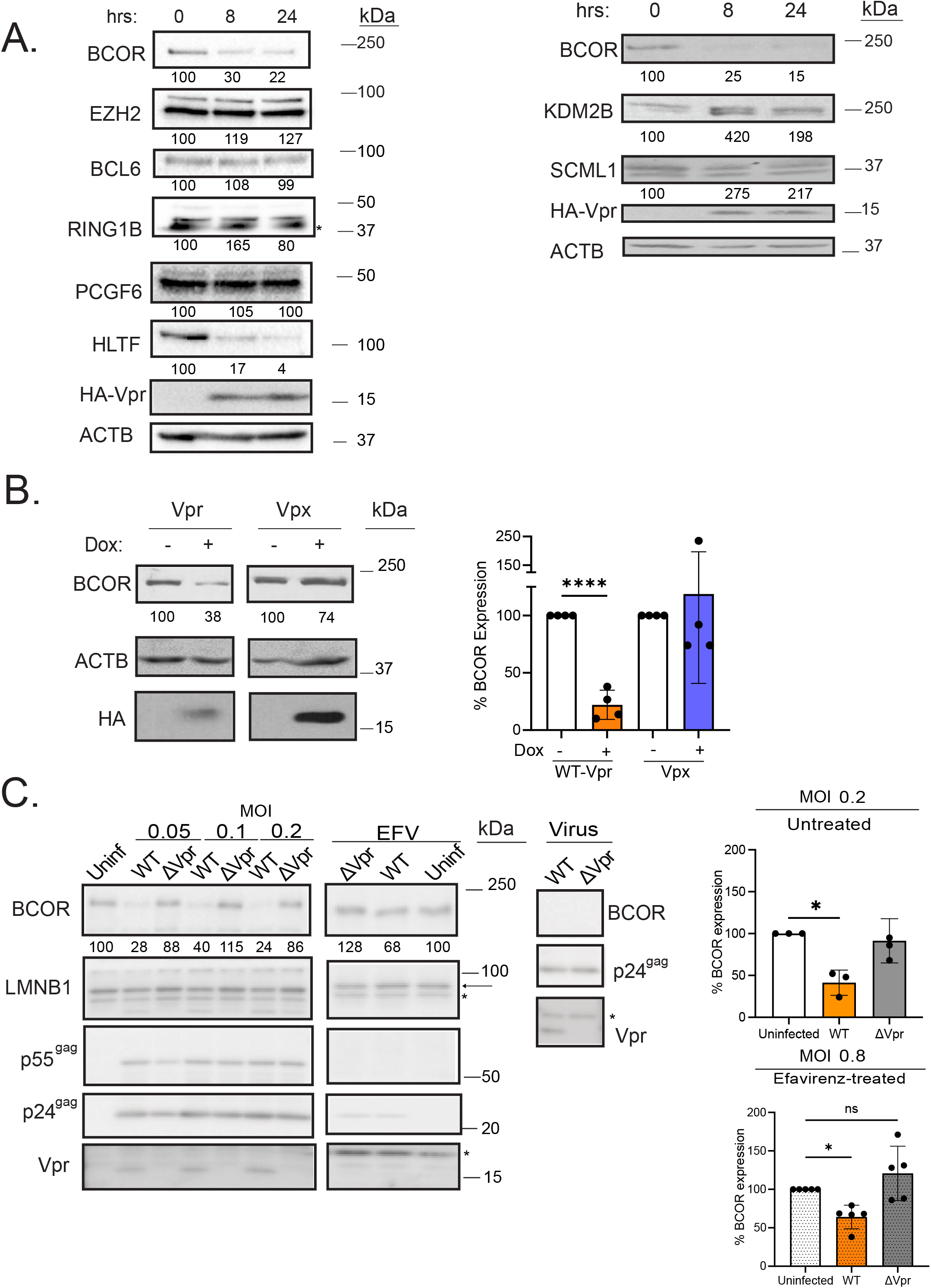
HIV-1 Vpr, but not HIV-2/SIVsmm Vpx, mediates BCOR depletion. (A) HIV-1 Vpr is sufficient to induce BCOR depletion. HeLa Flp-In T-Rex cells were treated with doxycycline for 8 or 24 h to induce HA-tagged HIV-1 Vpr expression. Cells were lysed and analyzed by immunoblotting for the indicated proteins. Protein bands were first normalized to the loading control, actin B (ACTB), and then to the uninduced control (0 h), which was set at 100%. Unspecific protein bands are indicated with an asterisk (*). Shown are representative immunoblots from three experiments. (B) The HIV-2/SIV_smm_ Vpx paralog is unable to induce BCOR depletion. Expression of HA-WT-Vpr or HA-Vpx (SIV_smm_) was induced for 24 h. Shown is a representative immunoblot panel (left panel with quantification of BCOR expression indicated below) and a summary graph analyzing the percent BCOR expression of four experiments (right panel). (C) Vpr induces BCOR depletion during HIV-1 infection of CD4^+^ T cells. MT4 cells were infected for 48h at the indicated MOIs with the GFP-reporter WT- or ΔVpr NL4-3 viruses (left panels) in the absence or presence of 5µM of the RT inhibitor Efavirenz (EFV). Cell lysates were analyzed by immunoblotting for the indicated proteins (quantification of BCOR expression is indicated below; LMNB1: lamin B1). The WT and ΔVpr NL4-3 virus used to initiate infection were also analyzed for the indicated proteins (right panel (virus)). Shown is a summary graph of untreated (upper right panel, n=3) or EFV-treated (lower right panel, n=5) infected cells at MOI of 0.2 and 0.8, respectively. The unpaired t-test was utilized in panel B while the ordinary one-way Anova test was used in panel C. Each dot on graphs represents an independent experiment. Unspecific protein signals are indicated with an asterisk (*). Uninf = uninfected. Data are represented as mean +/- SD.; *p< 0.5; ****p<0.0001.

Previous studies suggested that Vpx from HIV-2/SIV_smm_ (sooty mangabey) arose from a gene duplication of an ancestral Vpr when the SIV and HIV-2 groups diverged from a common ancestor (46). Both Vpr and Vpx are incorporated in virions and mediate the proteosomal degradation of specific host proteins, despite having very distinct biological activities (47, 48). Here, HeLa Flp-In T-Rex cells encoding SIV_smm_Vpx or HIV-1 Vpr were induced for 24 h to allow maximum expression of both proteins and then analyzed for BCOR expression. As shown in Figure 1B, BCOR depletion was observed with HIV-1 Vpr but not with SIV_smm_Vpx, implying that BCOR-depletion is not a function that HIV-1 Vpr shares with SIV_smm_ Vpx.

To assess whether the Vpr-mediated depletion of BCOR observed in our inducible system could be recapitulated in a *bona fide* infection setting, we infected the highly susceptible human T cell line, MT4, with a WT or ΔVpr GFP-expressing NL4.3 viral strain (49) (50) at different multiplicity of infection (MOI) and analyzed infected cell lysates by immunoblot. BCOR expression was consistently reduced in cells infected with WT virus compared to uninfected cells, whereas no significant reduction of BCOR was induced by the ΔVpr virus (Figure 1C; left panel and quantitation at 0.2 MOI on the right upper panel). To determine if virion-encapsidated Vpr was sufficient to induce BCOR depletion, MT4 were treated with the reverse transcriptase (RT) inhibitor Efavirenz (EFV) during viral adsorption with a pNL4.3 WT or ΔVpr GFP virus (Figure 1 C, virus panel). After 48h of continuous EFV treatment, we observed an average of 33% BCOR depletion in cells exposed to WT Vpr virus (Figure 1C; right panel (EFV) and quantitation at 0.8 MOI on the lower right panel) suggesting that virion-associated Vpr targets BCOR during the early phases of the infection prior to viral integration and *de novo* Vpr expression.

Taken together, our data indicate that BCOR, a core component of the ncPRC1.1 complex, is a putative target of Vpr during HIV-1 infection of CD4^+^T cells and further show that Vpr is sufficient to deplete BCOR protein levels.

### BCOR depletion and G2/M cell cycle arrest are two distinct functions of Vpr

The expression levels of many cellular proteins are regulated in a cell cycle-dependent manner and notably during the G2/M cell cycle arrest induced by HIV-1 Vpr (51) (23). To assess whether BCOR depletion is dependent from Vpr-mediated G2/M arrest, HeLa Flp- In T-Rex cells expressing no Vpr (parental cells), WT-Vpr or the well-established cell cycle arrest-impaired mutants R80A-Vpr and S79A-Vpr (52, 53) were analyzed for BCOR levels following doxycycline induction for up to 24 h. As shown in Figure 2A, we observed a time-dependent increase in the frequency of cells arrested at G2/M starting at 8 h post induction of WT Vpr and peaking at 16 h, while no cell cycle arrest was observed upon induction of R80A-Vpr. Interestingly BCOR had already been largely depleted by 8 h post doxycycline treatment (Figure 2B), suggesting that BCOR depletion does not result from the G2 cell cycle arrest induced by Vpr. In contrast, BCOR expression remained stable in the doxycycline-treated parental cells (Figure 2B) and, cell staining with a viability dye showed that the rate of cell mortality was comparable between parental and Vpr-expressing cells post doxycycline treatment (Figure S1A), further asserting the specificity of this Vpr phenotype. As expected, HLTF was also depleted to a comparable extent as BCOR upon Vpr-induced expression (Figure 2B; left and right panels). Using this system, it was clear that although Vpr derivatives, S79A and R80A, were unable to induce a G2/M cell cycle arrest (Figure 2A and Figure S1B), both mutants retain a substantial capacity to deplete BCOR and HLTF (Figure 2C; Figure S1C). Thus, these results support the notion that Vpr-dependent BCOR depletion is independent of the viral protein’s ability to induce an efficient G2/M cell cycle arrest.

**Figure 2.**
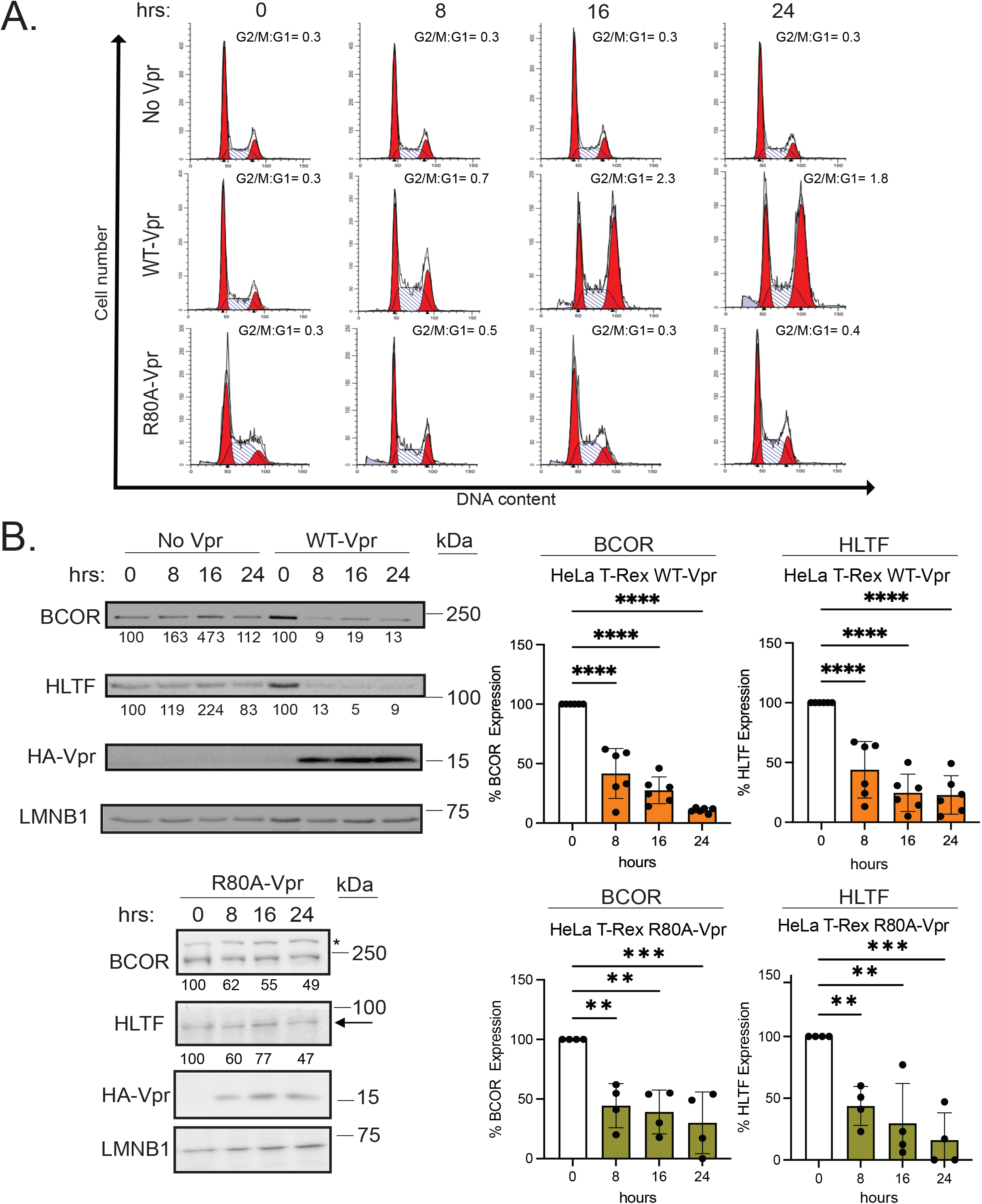
Vpr induces BCOR depletion independently of its G2/M cell cycle arrest activity. (A) Vpr induces a G2 arrest while the R80A Vpr mutant does not. HeLa Flp-In T-Rex cells expressing or not HA-WT-or HA-R80A-Vpr were treated with doxycycline for 8, 16, or 24 h and processed for cell cycle analysis by PI staining and flow cytometry. Shown is a representative cell cycle profile from one out of two experiments. Numbers over histograms represent the ratio of cells in G2/M over G1. (B) Cell cycle defective R80A Vpr mutant induces BCOR depletion. Cell samples processed as in panel A were also lysed and analyzed by western blot for the indicated proteins. Shown are representative blots (left upper and bottom panels) with protein band quantification assessed as described in Figure 1A. Right panels depict graphs summarizing the analysis of each experiment; HA-WT-Vpr n=6; HA-WT-R80A n=4. Unspecific protein bands are indicated with an asterisk (*). Error bars indicated +/- SD. Statistical analysis: Ordinary One-way Anova, **p<0.01, ***p<0.001, ****p<0.0001.

### BCOR depletion requires Vpr binding to DCAF1

Vpr depletes its targets by hijacking the CRL4^DCAF1^ ubiquitin ligase complex (13, 54). The Q65R mutation abrogates specifically the ability of Vpr to bind DCAF1 (16). Thus, to test if binding to DCAF1 is essential for Vpr-mediated BCOR depletion, the expression levels of BCOR were compared between WT-Vpr and the Q65R-Vpr mutant in the context of HeLa Flp-In T-Rex cells induced for 24h with doxycycline. Figure 3A (left panel) reveals that WT-Vpr but not Q65R-Vpr efficiently depletes BCOR (average 13% residual expression with WT Vpr *vs.* 118% with Q65R, left panel), suggesting that Vpr depletes BCOR via a process that requires binding to DCAF1. These findings were further validated by using siRNA that target DCAF1. Knocking-down DCAF1 abolished Vpr-dependent depletion of BCOR as well as of HLTF (Figure 3B). Overall, these results indicate that DCAF1 expression and binding are indeed critical for Vpr-mediated BCOR depletion, supporting the notion that the CRL4^DCAF1^ ubiquitin ligase complex is involved in this process.

**Figure 3.**
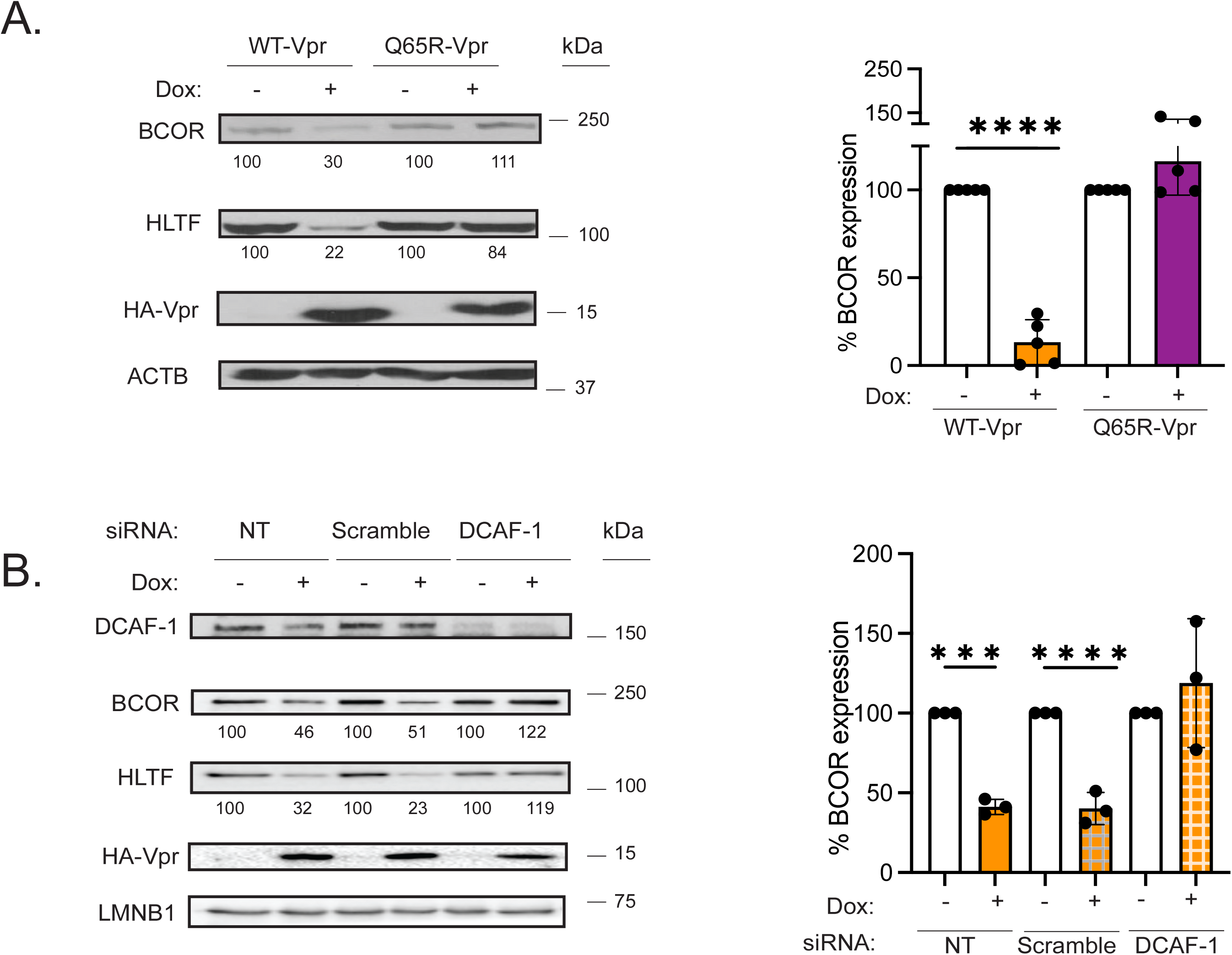
Vpr requires DCAF1 expression and recruitment to induce BCOR depletion. (A) The DCAF1-binding mutant, Q65R Vpr, is unable to induce BCOR depletion. HeLa Flp-In T-Rex cells were treated with doxycycline for 24 h to induce expression of HA-WT- or -HA-Q65R-Vpr and analyzed by Western blot for the indicated protein (left panel). BCOR and HLTF protein bands quantification are indicated below the blots. Shown is a graph summarizing the quantitation analysis of BCOR expression (right panel, n=5). (B) HeLa Flp-In T-Rex cells were transfected or not with Scramble or DCAF1-targeting siRNA for 24h and then treated with doxycycline for 24h. Cell lysates were analyzed by Western Blot for the indicated protein (left panel with quantification of BCOR and HLTF expression indicated below). Shown is a graph summarizing the quantitative analysis of BCOR expression in three experiments (right panel). For all panels, error bars indicate +/-SD, unpaired t-test was used, ***p< 0.001, ****p<0.0001.

### Vpr-dependent BCOR depletion requires functional proteasome and neddylation machineries

The hijacking of the CRL4^DCAF1^ ubiquitin ligase complex ultimately leads to the polyubiquitination of Vpr target proteins and their depletion via the proteasome (13, 54).To assess whether a functional proteasome was required for Vpr-dependent BCOR depletion, WT-Vpr HeLa Flp-In T-Rex cells were treated with DMSO, the proteasome inhibitor MG132 (10 µM) or the endolysosomal inhibitor chloroquine (100 µM) 1 h post doxycycline treatment and then analyzed for BCOR expression 7h later (Figure 4A). While BCOR was efficiently depleted in the DMSO control (approximately 38% average residual expression, left panel), and in the presence of chloroquine (approximately 28% average residual expression, left panel), the depletion was completely inhibited when cells were exposed to MG132 indicating the involvement of the proteasome, but not endolysosomal proteases, in Vpr-mediated BCOR depletion (Figure 4A).

**Figure 4.**
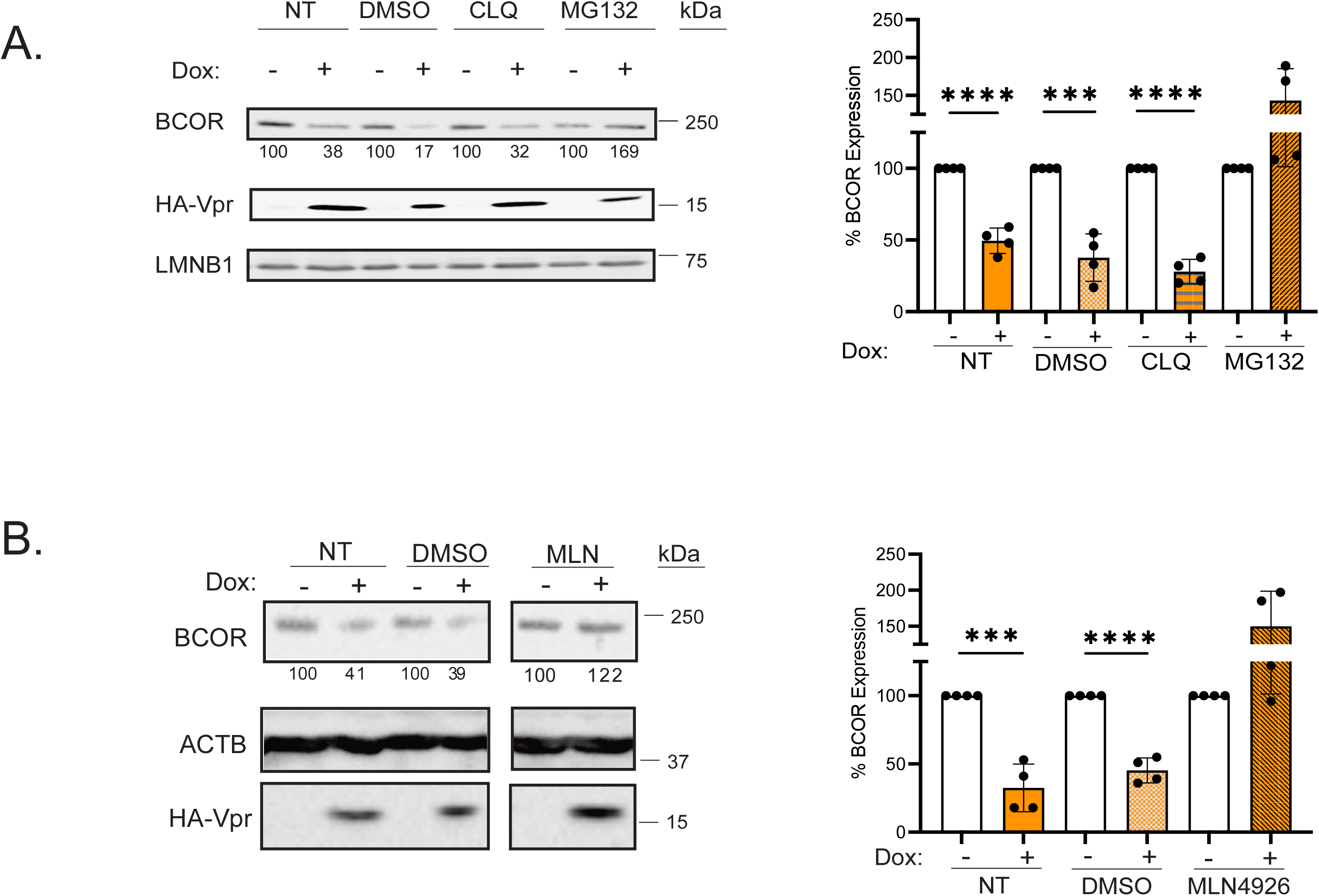
BCOR depletion by Vpr requires functional proteasome and neddylation pathways. (A) Vpr-mediated BCOR depletion is inhibited by the proteasome inhibitor MG132. HA-WT-Vpr expression was induced for a total of 8 h. An hour post doxycycline treatment, cells were either left untreated (NT), treated with DMSO, Chloroquine (CLQ), or MG132. Cell lysates were then analyzed by immunoblotting for the indicated proteins (left panel with BCOR expression indicated below). Shown is the quantitative analysis of BCOR expression from four experiments (right panel). (B) Vpr-mediated BCOR depletion is inhibited by a neddylation inhibitor. HA-WT-Vpr expression was induced as described above; cells were then either left untreated (NT), treated with DMSO or treated with MLN4926 (MLN). Cell lysates were analyzed for the indicated proteins by immunoblotting (left panel with quantification of BCOR expression indicated below). Shown is a graph summarizing the quantitative analysis of BCOR expression in 4 experiments (right panel). For all panels, data are represented as mean +/- SD. Unpaired t-test was utilized ***p< 0.001, ****p<0.0001.

The CRL4^DCAF1^ complex assembles with the scaffold protein, Cullin4A (55). Neddylation of Cullin proteins by the enzyme neural precursor cell expressed developmentally down-regulated protein 8 (NEDD8) is critical for the ubiquitin E3 ligase activity of the complex (56). To evaluate if the proteasome-dependent degradation of BCOR requires neddylation, HeLa Flp-In T-Rex HA-WT-Vpr cells were treated 1 h post doxycycline treatment with the NEDD8 inhibitor, MLN4924 (10 µM), and then analyzed for BCOR expression 7h later (Figure 4B). While BCOR was degraded when WT-Vpr-induced cells were treated with DMSO as control (45% average residual expression, left panel), treatment with MLN3924 completely inhibited BCOR degradation (Figure 4B). These results indicate that Vpr-mediated BCOR degradation requires a functional NEDD8 and further support the notion that the CRL4^DCAF1^ E3 ligase complex is essential for this activity.

### BCOR contributes to HIV-1 gene silencing during viral latency

BCOR is mainly known to function in the repression of gene expression (57). We hypothesized that HIV-1 evolved to target BCOR for proteasomal degradation to counter PRC-dependent provirus transcription silencing, therefore allowing optimal viral expression. To gain insight as to whether BCOR is directly involved in the transcriptional repression of HIV-1, we took advantage of the CD4^+^T latent cell line Jurkat E4 model of HIV latency, which carries at least one copy of an integrated GFP-marked Gag- and Vif-, Vpr-, Nef-defective provirus (58) (Figure 5A), and expresses detectable BCOR protein (Figure S2A). These cells were selected for their ability to silence the integrated HIV-1 provirus and as such represent a relevant cellular model where epigenetic repression is exerted on the integrated provirus (41). In this context, ChIP-qPCR revealed the occupancy of BCOR and the associated ncPRC1.1 component, PCGF1 on the LTR as well as the presence of the distinctive silencing histone mark H2AK119ub using primers covering nucleosomes (Nuc0 and Nuc1) and the transcription starting site (TSS) (Figure 5B and Figure S2B). In addition, as previously reported (36), the well-studied PRC2 methyltransferase, EZH2, was found recruited to the LTR in these cells as well as the repressive chromatin modification it catalyzes on histone 3, H3K27me3 (Figure 5B), suggesting that the BCOR-containing ncPRC1.1 contributes along with PRC2 to the silencing of the integrated HIV in the Jurkat E4 model of latency.

**Figure 5.**
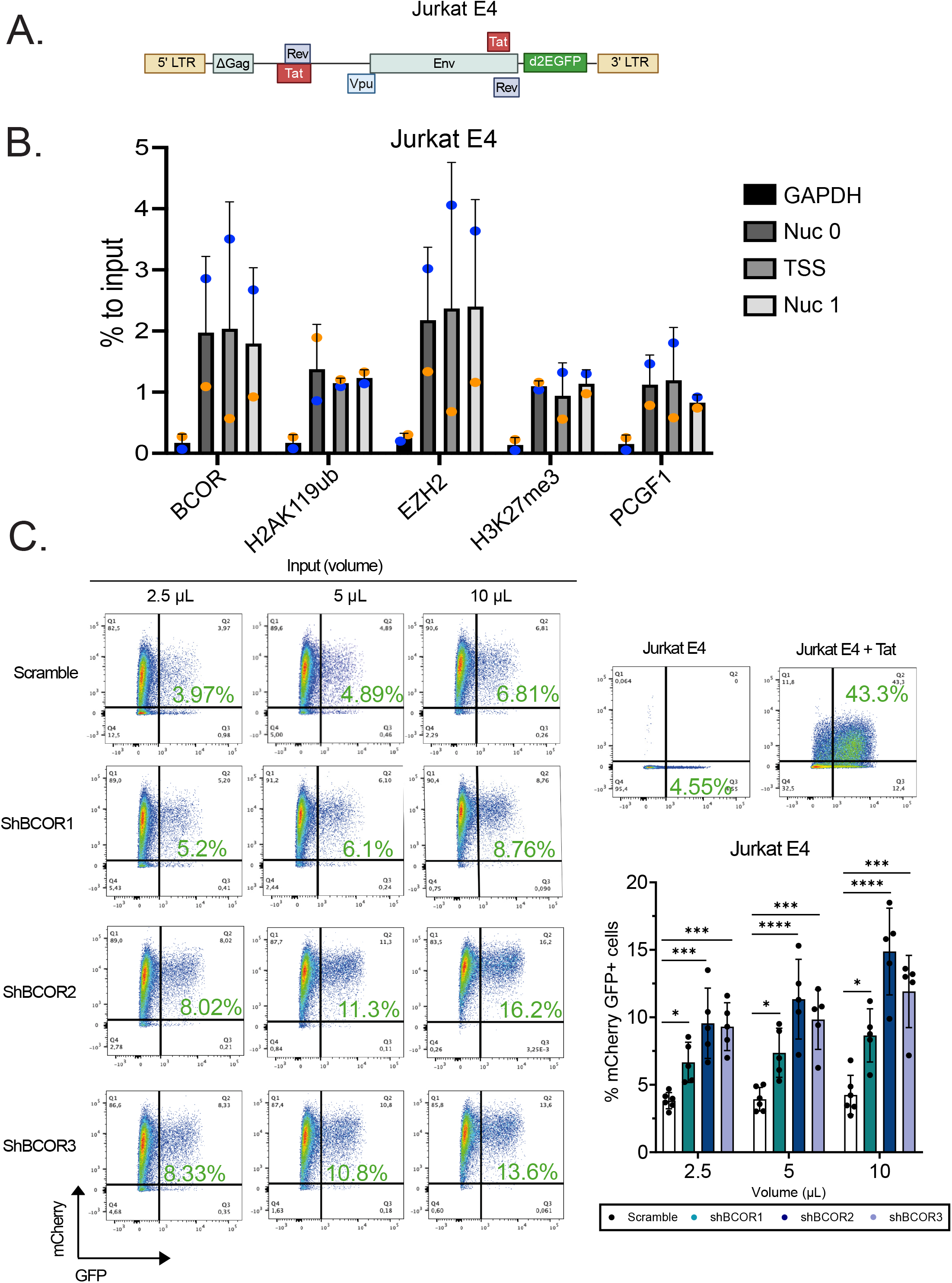
BCOR contributes to HIV-1 transcriptional silencing. (A) Schematic representing HIV-1 provirus integrated in Jurkat E4 CD4^+^ T cells, which express a GFP with a short half-life (d2EGFP). (B) Analysis of the occupancy of BCOR, EZH2, PCGF1 and histones with silencing marks H2AK119ub or H3K27me3 on the HIV-1 LTR. Histograms representing percentage of input recruited to Nuc 0, transcriptional starting site (TSS) or, Nuc 1 from two independent color-coded experiments. GAPDH promoter sequence was used as a negative control genomic region. (C) shRNA-directed BCOR depletion enhances expression of latent HIV-1. Jurkat E4 were transduced with mCherry-reporter lentivirus expressing either scramble or BCOR-targeting shRNAs (#1, #2, or #3) at the indicated input doses or the viral protein Tat at 2.5 μL of lentiviral vector. Shown is a representative analysis of GFP expression in the indicated transduced Jurkat E4 samples (mCherry^+^) by flow cytometry (left and upper right panels). A graph summarizing the analysis of five independent BCOR depletion experiments with the indicated shRNA is also shown (bottom right panel). Data are represented as mean +/- SD. Statistical analysis: Ordinary one-way Anova test *p< 0.5, ***p<0.001, ****p<0.0001.

To evaluate the contribution of BCOR to LTR silencing, Jurkat E4 cells were transduced with mCherry-reporter lentiviruses expressing either scrambled or BCOR-targeting shRNAs (#1, #2 or #3) (Figure S2C) and analyzed for both HIV-1 LTR-driven GFP expression and BCOR expression. Transduction of Jurkat E4 cells with all three BCOR-targeting shRNAs, but not the scramble control, led to a significant reactivation of LTR-directed GFP expression in transduced cells, with shRNA #2 and #3 noticeably inducing a two-fold increase in the frequency of mCherry^+^ GFP^+^ (Figure 5C). This upregulation of LTR-directed GFP expression was linked to a ∼ 40 to 50% knockdown of BCOR expression across the total cell population in the presence of shRNA #1-3 relative to cells transduced with the scramble shRNA control (Figure S2D). Interestingly, transduction of a lentivirus expressing Tat increased the frequency of mCherry^+^GFP^+^ cells by approximately 10-fold on average (Figure 5C), indicating that a larger population of integrated proviruses were reactivable by Tat overexpression.

A significant occupancy of the three distinct LTR regions by BCOR, PCGF1, H2AK119ub, EZH2 and H3K27me^3^ was also detected in the Jurkat 2D10 HIV latency model, which contains an integrated GFP-marked HIV-1 provirus similar to that of Jurkat E4 except that it encodes an attenuated form of Tat (H13L) (58) (Figure S3A and Figure S3B). Consistent with the results observed in Jurkat E4, transduction of shRNA #2 in 2D10 cells induced a two-fold increase in the frequency of mCherry^+^GFP^+^ cells (Figure S3C), although the frequency of mCherry^+^GFP^+^ cells was overall lower in this model cell line most likely because of the altered Tat function encoded by the integrated provirus.

Taken together, these results, indicate that the ncPRC1.1 components, BCOR and PCGF1, occupy the HIV-1 LTR along with their distinctive silencing mark, H2AK119ub, in two CD4+ T cell models of HIV-1 latency and further reveal that knock-down of BCOR enhance LTR-directed gene expression thus implying that BCOR is contributing to the PRC-dependent repression of the HIV-1 LTR-directed transcription.

### Vpr-mediated BCOR depletion reactivates expression of integrated latent HIV-1

Our findings thus far reveal that Vpr degrades BCOR and that BCOR contributes to repression of HIV-1 LTR-driven expression in cellular models of HIV-1 latency. Consequently, we postulated that Vpr modulation of BCOR expression would ultimately lead to reactivation of integrated proviruses in latently infected cells. To this end, Jurkat E4 cells were transduced for 24 h with a lentivirus expressing a polycistronic mRNA encoding either WT-Vpr, Q65R-Vpr or Tat under the control of the EF1α promoter and co-expressing a mCherry reporter gene through an IRES sequence (Figure 6A). To assess if virion-associated Vpr is sufficient to induce viral reactivation, transductions were also performed in condition where cells were exposed to Efavirenz (EFV) during and after adsorption with Vpr-expressing lentiviral vectors for 24 h. Cells were then analyzed by flow cytometry to determine transduction efficiency (mCherry^+^) and provirus reactivation (GFP^+^) (Figure 6B). While Jurkat E4 transduced with lentivirus encoding WT-Vpr, displayed significant HIV-1 reactivation levels (average of 2.8-fold), cells transduced with the Q65R-Vpr counterpart didn’t exhibit any increase in the frequency of mCherry^+^/GFP^+^ positive cells relative to the negative control (Figure 6B, middle left panel and quantification in the right upper panel). As shown in Figure 6C, the extent of HIV reactivation was linked to Vpr-mediated BCOR depletion (compare BCOR expression in the presence of WT Vpr and the Q65R Vpr mutant). EFV treatment, which allows to evaluate the effect of virion-associated Vpr, revealed an average of 2.5-fold of reactivation of the HIV-1 LTR by WT-Vpr but not by Q65R-Vpr (Figure 6B, lower left panel and quantification in the right panel). The degree of Vpr-mediated reactivation of integrated HIV-1 in condition of EFV treatment was similar to the untreated conditions, and both conditions were linked to a comparable degree of BCOR depletion (Figure 6C). Overall, these results suggest that Vpr promotes reactivation of latent HIV proviruses by targeting BCOR to proteasomal degradation through a process that involves the recruitment of the CRL4^DCAF1^ E3 ligase. The fact that virion-associated Vpr has the capacity to reactivate latent integrated HIV-1 proviruses, suggests that Vpr packaged into incoming viral particles could target BCOR-mediated silencing processes during the early stages of infection prior or/and during proviral DNA integration, thus allowing efficient viral expression from integrated proviruses.

**Figure 6.**
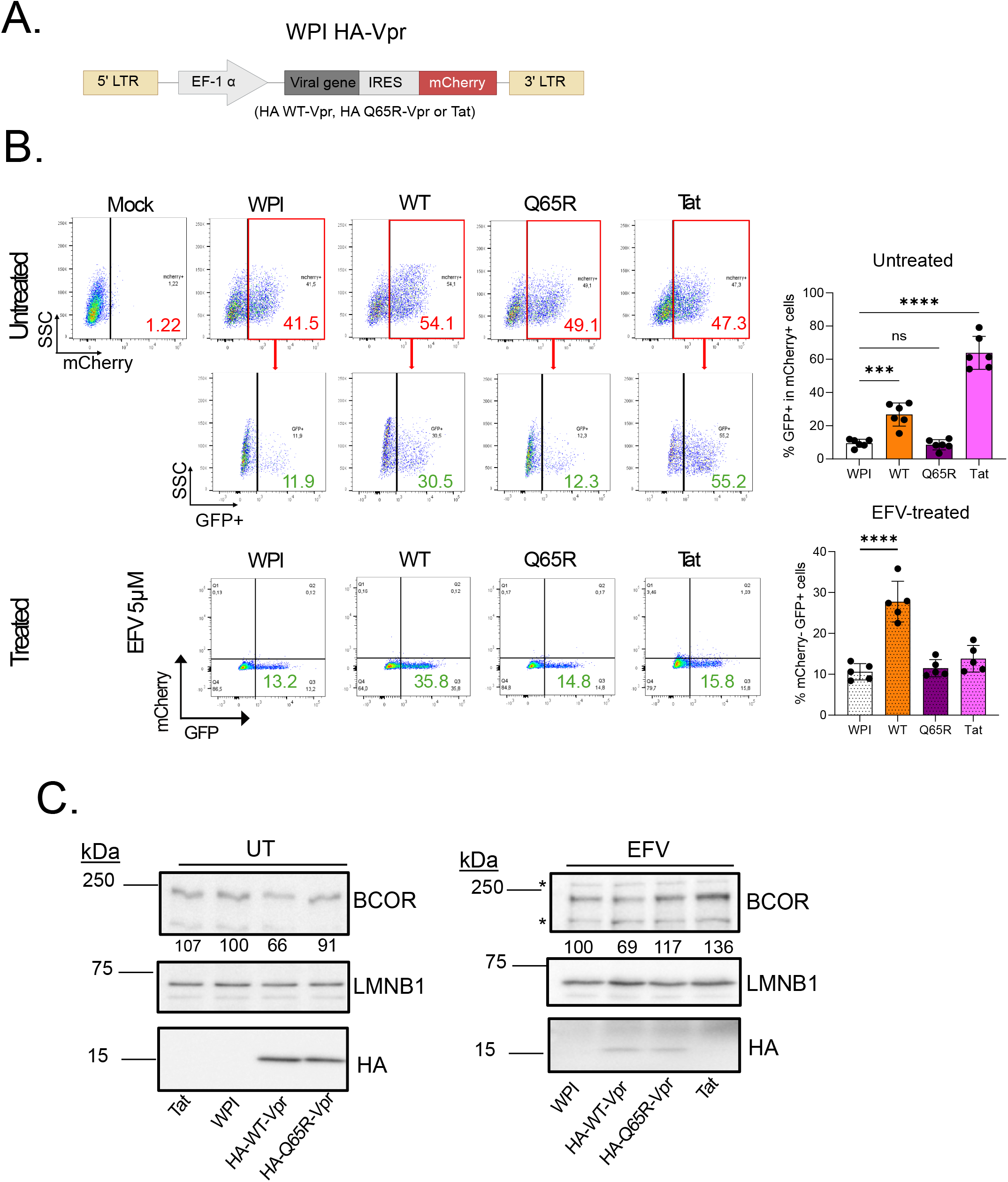
Vpr reactivates latently infected cells. (A) Schematic representation of the lentiviral vector expressing HA-Vpr-, Q65R HA-Vpr- or Tat-IRES-mCherry. (B) Transduction of Vpr reactivates latent HIV-1 expression. Jurkat E4 cells were transduced with 2.5 μl of the mCherry-reporter lentivirus expressing WT-, Q65R-, or a control vector in the absence or the presence of 5µM EFV. Representative analysis of the % of GFP^+^ cells in the transduced mCherry^+^ population (left panel). Numbers in the right quadrant represent the percentage of transduced mCherry^+^ cells in the different lentiviral transduction conditions (top panel; red). Numbers in the bottom panels represent the percentage of reactivated GFP^+^ cells (green) in the transduced mCherry^+^. Shown is a summary graph of the average percentage of GFP^+^ cells within mCherry^+^ cells in non-treated (n=6) (upper right panel) or for EFV-treated conditions (n=5) (lower right panel). Data are represented as mean +/- SD. Statistical analysis: Ordinary one-way Anova test, ***p<0.001, ****p<0.0001, ns, not significant. (C) Activation of HIV expression in Jurkat E4 cells by Vpr is linked to BCOR depletion. Cell samples were also collected and analyzed by western blot for the indicated proteins. Shown is a representative of three independent experiments (UT= untreated). Unspecific protein bands are indicated with an asterisk (*).

## DISCUSSION

HIV-1 transcription is known to be sensitive to the chromosomal environment at the site of integration with proviruses integrated into centromeric heterochromatin displaying extensive epigenetic silencing marks and undetectable levels of basal transcription (2). Epigenetic factors silencing HIV-1 gene expression are of particular interest since they contribute to restricting virus replication while at the same time influencing the establishment and maintenance of persistent latent reservoirs capable of reseeding virus production after cessation of antiviral therapy. Recent evidence indicates that primate lentiviruses have evolved mechanisms to counter transcriptional silencing at pre- and post-integration stages suggesting that these processes are regulated during infection (11, 59, 60). Here, we provide evidence indicating that the HIV-1 Vpr accessory protein, but not its paralog Vpx, encoded by lentiviruses from the SIVsmm/HIV-2 lineages, mediates the degradation of BCOR, a component of the PRC system known to be involved in silencing HIV-1 expression (37, 60) (Figure 1). Vpr being the accessory protein most abundantly packaged into nascent HIV-1 particles (61), we further show that incoming virion-associated Vpr can recapitulate the degradation of BCOR observed during virus infection or exogenous Vpr expression, indicating that HIV-1 Vpr is necessary and sufficient to trigger BCOR degradation (Figure 1).These results are consistent with previous unbiased proteomic analysis revealing that HIV-1 Vpr is largely responsible for a global cellular remodeling, involving targeting and depletion of multiple mostly nuclear cellular proteins, among which BCOR was identified (23). Interestingly, quantitative proteomics analysis revealed that profound depletion of BCOR was observed with Vpr variants from the HIV-1/SIVcpz lineage, including Vpr derived from primary virus belonging to group M (clade C) and group O, but not with Vpr nor with Vpx from SIVsmm/HIV-2 lineage. These findings suggest that Vpr-mediated BCOR depletion is a conserved feature unique to viruses from the HIV-1/SIVcpz lineage and further support the notion that SIVcpz, the precursor of HIV-1, has evolved to mediate this Vpr-associated activity prior to the cross-species transmission that led to the emergence of HIV-1. This specific difference between HIV-1 and HIV-2 Vpr might be relevant *in vivo* since it was proposed that the limited global proteome remodeling induced by HIV-2 Vpr and Vpx may explain in part the reduced pathogenicity of HIV-2 in human over HIV-1 (23) .

BCOR is the first PRC-associated component known to be countered by HIV-1, specifically through Vpr hijacking of the CRL4^DCAF1^ ubiquitin ligase and targeting of the co-repressor for proteasomal degradation. None of the other PRC components identified in our previous BioID analysis (24) as putative Vpr interacting partners, including EZH2, Ring1B, PCGF6, KDM2B and SCML1, were depleted by Vpr (Figure 1). The degradation of BCOR by Vpr, which requires the expression and recruitment of DCAF1 (Figure 3) as well as the neddylation of Cullin and an active proteasome (Figure 4), is shown to be completely independent from its ability to induce G2/M cell cycle arrest or cell death (Figure 2). Indeed, well-characterized mutants of Vpr, VprR80A and Vpr 79A, which are G2-arrest-defective but competent for recruiting DCAF1 (20, 23, 24) retain the capacity to mediate the degradation of BCOR. This finding is also supported by the observation that HIV-2 Vpr is capable of inducing a G2 arrest (62) but is unable to deplete BCOR (23).Together, these results imply that PRC-mediated transcriptional silencing has likely exerted some evolutionary pressure on HIV-1. Previous studies have shown that shRNA-mediated depletion of the histone methyltransferase component of PRC2, EZH2, or PCGF4 found either in cPRC1 or ncPRC, alone were sufficient to reactivate latently infected cells (37–41, 58, 63–65). Our results reveal that shRNA-directed BCOR depletion results in activation of HIV-1 gene expression in the Jurkat E4 and 2D10 T cell models of HIV-1 latency (figure 5). This activation of HIV-1 expression was phenocopied by lentiviral mediated transduction of Vpr in Jurkat E4 and was linked to a depletion of BCOR (Figure 6). Indeed, in both Jurkat E4 and 2D10 T cells, we show not only the occupancy of EZH2 and its distinctive tri-methylation repressive marks on H3K27 on the LTR as was previously reported (36, 38, 41) but also the occupancy of the BCOR and PCGF1 components of the ncPRC1.1 and its distinctive ubiquitination marks on H2AK119 (Figure 5 and Figure S3). The fact that independent depletion of BCOR, EZH2 and PCGF4 is sufficient to reactivate latently infected cells strengthens the possibility that each type of PRC complexes, namely ncPRC1, cPRC1, and PRC2 is required for the stable long-term silencing of HIV-1 expression. Indeed, it was reported that transcriptional silencing by the PRC system can function in a sequential manner whereby ncPRC1 recruit and/or stabilize PRC2 and stall RNA polymerase II via ubiquitination of histone 2A at lysine 119 (H2AK119ub), then PRC2 tri-methylates histone 3 at lysine 27 (H3K27me3) to recruit cPRC1, which subsequently compacts the chromatin (29–32) . Our results support the notion that HIV-1 Vpr-mediated degradation of the ncPRC1.1 core component BCOR initiates a cascade of destabilization starting with ncPRC1.1 that subsequently impairs the functional roles of the other PRC complexes, thus countering PRC-mediated silencing of HIV-1 transcription.

Overall, our results indicate that HIV-1 Vpr drives BCOR degradation to counter PRC-dependent transcriptional silencing of theHIV-1 integrated provirus. The fact that BCOR-depletion can be induced by virion-associated Vpr suggests that this PRC counteraction may occur at early stages of infection prior or during provirus integration, in order to create a chromosomal environment conducive to early viral transcription before expression of the Tat transactivator. These findings provide insights not only on the PRC system complexes that are sequentially involved in repressing HIV-1 expression and maintaining latency in CD4^+^ T cells but also on how HIV-1 evades transcriptional silencing to achieve optimal expression and replication. Hence, the present study underscores the importance of further investigating host factors that repress proviral expression to develop effective strategies to purge the latent HIV-1 reservoir.

## MATERIALS AND METHODS

### Cell culture

The generation of HeLa Flp-In T-Rex cell lines carrying one copy of HA-tagged HIV-1_Bru_-WT, -Q65R, -S79A or -R80A-Vpr was described previously using the parental cell line HeLa Flp-In T-Rex (RRID:CVCL_C4ET) (24).

HeLa Flp-In T-Rex HA-Vpx cell line was generated in this study using HeLa Flp-In T-Rex parental cell line. HA-Vpx was amplified by PCR amplification from a pUC57-HAVpx_smm_ encoding plasmid (66), using the primers described in Table 1. Cloning and stable cell line generation was done as previously described (24).

The BCOR knockdown (KO) 293T cell line was generated by transfecting HEK 293T (RRID: CVCL_0063; ATCC) with the pSpCas9(BB)-2A-Puro V 2.0 system encoding a gRNA targeting *BCOR* exon 6. Cloning of the gRNA was performed using the primers listed in Table 1 as described (67). Cells were transfected using lipofectamine 2000 following manufacturer instructions and selected 48 hours post-transfection with puromycin (1mg/mL).

HeLa Flp-In T-Rex, HEK293T and BCOR KO 293T cells were cultured in DMEM media supplemented with 10% FBS. HeLa Flp-In T-Rex cell lines were maintained in the presence of hygromycin (50 µg/ml) and doxycycline (0.5 µg/ml) was used to induce HA-tagged Vpr, Vpr mutants, or Vpx expression. For protein degradation assays, cells were treated with either 10 µM MG132, 10 µM MLN4924, 100 µM chloroquine or DMSO, 1 hour post doxycycline induction.

MT4 (RRID: CVCL_2632; NIH AIDS Reagent Program), Jurkat E6.1 (RRID: CVCL_0367; ATCC), Jurkat E4 and 2D10 cells (provided by Dr. Jonathan Karn; (58) ) were all grown in RPMI media containing 10% FBS. All cell lines were regularly tested and verified to be free of mycoplasma.

### Construction of mCherry-expressing lentiviral vectors

pHRSiren plasmid expressing mCherry was generated by digestion of the pHRSiren plasmid with PacI and KpnI followed by ligation of the mCherry reporter downstream of the spleen focus-forming virus (SFFV) promoter. Scramble or BCOR-targeting shRNA (see table 1 for sequences) containing the digested BamH1 and EcoR1 restriction sites were annealed, gel purified and then ligated into the BamH1/EcoR1 digested mCherry-expressing pHRSiren plasmid.

To replace GFP with mCherry in the Vpr-expressing WPI lentiviral plasmids (68), HA-Vpr-expressing vectors were digested with BmgBI and BstBI and purified to remove the GFP gene. mCherry from pHR-Siren was cloned using the previously mentioned restriction sites to generate pWPI-derived HA-WT-Vpr-IRES-mCherry, HA-Q65R-Vpr-IRES-mCherry or Control-IRES-mCherry vectors. Tat-IRES-mCherry plasmid was generated by overlapping PCR of the Tat cDNA contained in a SVCMV-Tat expression plasmid (69) and the IRES mCherry sequence from HA-WT-Vpr-IRES-mCherry. The resulting PCR product was cloned into PmeI and BstBI sites of the HA-WT-Vpr-IRES-mCherry plasmid. See Table 1 for primer sequences. All constructs were confirmed by Sanger sequencing.

### Production of HIV-1 and lentiviral-based vectors

Virus and lentiviral vectors were generated by transfection of proviral constructs into HEK293T cells using the calcium phosphate method as previously described (24). Briefly, for production of GFP-marked NL4-3 WT and ΔVpr (50), 7×10^6^ HEK293T cells were plated onto 15 cm plates and transfected with 30 µg of proviral DNA constructs 24h after. For production of HA-Q65R-, HA-WT-Vpr-,Tat-IRES-mCherry and shRNAs lentiviral vectors, 20 µg of pWPI-derived or shRNA -expressing lentiviral vector, 20 µg of PAX2, and 7 µg of VSV-G were co-transfected into HEK293T cells (68). Media was replaced 12-16 h later, and 48-60 h post transfection, virus-containing supernatant was concentrated by ultra-centrifugation (35,000 RPM at 4°C for 2 h) onto a 20% sucrose cushion. Concentrated virus pellets were resuspended in 300 µl of media. Multiplicity of infection (MOI) was determined by infecting MT4 cells with increasing volume of infectious virus and calculating the number of infectious particles per microliter.

### Infections and transductions

MT4 cells were infected at the indicated MOI in round-bottom 96-well plates (1 ×10^6^ cells /100 µl). After 4 h of viral adsorption, cells were washed and placed in a 6-well plate for 2 days in a final volume of 3 ml. For transduction, Jurkat E4 or 2D10 cells lines (5 ×10^5^ cells) were plated and treated with 2.5, 5 or 10 µl of concentrated lentiviral vector expressing shRNA or HA-Q65R, HA-WT-Vpr, control or Tat. Cells transduced with shRNA-encoding vectors or Tat-IRES-mCherry were collected 96 h postransduction while cells transduced with HA-Q65R-, HA-WT-Vpr-, Control-IRES-mCherry or with Tat-IRES-mCherry vectors were collected after 24 h for flow cytometry analysis and/or immunoblot. For Efavirenz-related experiments, cells were treated with 5 µM Efavirenz 16 hours prior to infection or transduction. After viral adsorption, fresh Efavirenz was added and treatment continued for 24 to 48 hours depending on the experiment.

### Western blotting

Cell pellets were lysed with RIPA buffer (0.2% SDS, 1% deoxycholate, 2% NP-40, 5% glycerol, 150 mM NaCl, 25 mM Tris (pH 7.4), 1mM EDTA, and complete protease inhibitors) on ice for 45-60 min. Lysates were then clarified by centrifugation (10,000 RPM at 4°C for 10min). Total proteins from cell lysates were quantified using the BioRad protein assay (BioRad; 5000111). Up to 25-50 µg of cell extracts were resolved on (6%-15%) poly acrylamide gels and transferred to PVDF membranes. Anti-BCOR (Proteintech; 12107-01), anti-human DCAF1 (Bethyl; A301-887A), anti-human HLTF (Bethyl; A300-230-M), anti-human Lamin B1 (Abcam; ab6048), anti-human β-Actin (Abcam; ab49900), anti-HA (Biolegend; 901513), anti-human KDM2B (Millipore; 1710264), anti-human BCL6 (Cell Signaling; 14895S), anti-human EZH2 (Cell Signaling; 5246S), anti-human RING1B (Abcam; ab181140), anti-human RING1A (Abcam; ab175149), anti-human PCGF6 (Abcam; ab192395), anti-Gag p24 (ME Biotech Services; 31-90-25), anti-human SCML1 (Santa Cruz; SC135622), anti-Vpr R3.7 (Cohen et al., 1988), goat anti-rabbit-HRP (Abcam; ab205718) and goat anti-mouse-HRP (Abcam, ab205719) were used to detect target proteins. Membranes were exposed using a Bio-Rad Chemidoc MP imaging system. Densitometric quantification of protein bands was performed using Image J (v1.52).

### FACS staining and analysis

To analyze the cell cycle profile, cells were first resuspended in 100 µl of cold PBS and 900 µL of 70% ethanol was added prior to an incubation at 4°C. After washing with PBS, cells were exposed to RNAse (200 µg/ml) for 1 h at 37°C. Samples were then analyzed immediately after propidium iodide (PI) staining (final concentration, 30 µg/ml). All FACS analysis were performed on a BD Biosciences LSR Fortessa using the DIVA software (BD Bioscience). Data was analyzed using FlowJo (BD Life Sciences). Cell cycle analysis was performed using the ModFit mathematical model (Verity Software) to quantify the fraction of cells in G1 and G2/M phases.

To assess cell viability, cells were collected by centrifugation, stained with Fixable Zombie Aqua Live/Dead dye (Biolegend; 423101) for 15 minutes at room temperature and diluted 1:1000 in PBS buffer. Cells were washed twice with PBS 5% FBS followed by fixation with 2% paraformaldehyde (PFA).

### siRNA transfection

HeLa Flp-In T-Rex WT-Vpr cells (1.5×10^6^) were plated onto a 10-cm petri-dish and then transfected the next day with scramble or DCAF1-targeting siRNA at a final concentration of 500 nM using the Lipofectamine RNAi kit (Thermofisher; 13778075). Briefly, 15 µl of lipofectamine was diluted in 450 µl of Optimem media and siRNA was diluted in 450 µl of Optimem. Diluted siRNA and lipofectamine were then mixed, vortexed, and incubated for 5 min at room temperature in the dark before addition to plated cells. Media was changed 5 h post-transfection. Doxycycline was added 24 h post-transfection to induce Vpr expression. Cells were harvested for analysis 24 h post doxycycline treatment.

### ChIP-qPCR

For Jurkat E4, 1×10^7^ cells were used per immunoprecipitation. Cells were washed twice with PBS and treated with 2mM DSG (Thermo Scientific; 20593) crosslinker for 20 min.

Cells were washed 3 times with PBS and crosslinked with a final concentration of 1% formaldehyde for 10 min on a rocking platform at room temperature. Crosslinking was stopped by adding glycine at a final concentration of 125 mM and rocking for an additional 5 min. Cells were then washed twice at 4°C and centrifuged at 700 x *g* for 5 min with cold PBS. Cell pellets were frozen at -80°C. Dry cell pellets were lysed and chromatin was fragmented according to the Pierce Agarose ChIP kit (Thermo Scientific; 26156) manufacturer’s instructions. Lysate was precleared with 10 µL of A/G agarose beads overnight. Cleared samples were immunoprecipitated overnight with 2 µg of anti-BCOR (Proteintech; 12107-01), anti-human EZH2 (AC22) (Active Motif; 39076), anti-H3K27me3 (Abcam; ab6002), anti-H2AK119ub (Cell Signaling; 8240S) or anti-PCGF1 (AB_2716800, kindly provided by Dr. Vivian Bardwell (University of Minnesota)). 10% volume of the chromatin was used as input. After 2 h of incubation with A/G agarose beads, samples were washed for 15 min with buffers contained in the CHIP kit. Eluted samples were decrosslinked overnight at 65°C in presence of proteinase K.

For 2D10 experiments, 1×10^7^cells were crosslinked with 1% formaldehyde for 8 min on a rocker platform. Cross-linking was stopped by adding glycine at a final concentration of 125 mM and rocking for an additional 5 min. Cells were then washed 2 times with cold PBS. Cell pellets were frozen at -80°C. Dry pellets were lysed with RIPA buffer (0.1% SDS) for 1 h on ice and then sonicated with a Covaris sonicator (600 sec, a duty factor of 10, 140 peak incident power, and 200 burst count per sample). Lysates were precleared for 1 h and immunoprecipitated with 4-3 µg of antibody overnight in the presence of Dynabeads. 2% volume of the chromatin was used as input. Samples were washed once with RIPA buffer for 10 minutes followed by 3 washes with ChIP wash buffer (0.5M LiCl). Eluted samples were decrosslinked overnight at 65°C in presence of proteinase K.

Each sample was analyzed by qPCR using the SYBR-Green kit and previously published primers (36) numbered with respect to the transcriptional starting site of pHR LTR and targeting Nuc 0 (-390,-283), the transcription start site TSS (-116,+4) and Nuc1 (+30, +134) to assess the amount of chromatin immunoprecipitated by the antibodies. See Table 1 for sequence of primers. A final concentration of 0.5µM of each primer was mixed with 5 µl of SYBR-Green (Applied Biosystems) in final 10 µl of reaction. The qPCR was allowed to run for 40 cycles. Signal was normalized to the percentage of input method.

## QUANTIFICATION AND STATISTICAL ANALYSIS

Quantification of Western blots band signals was performed using Image J software. Treated or induced samples were first normalized to their loading control (ACTB or LMNB1) and then, to their corresponding untreated or uninduced controls, which was set at 100%, in all cases. Statistical analysis was performed using GraphPad Prism Version 8. Each dot represents an independent experiment, and error bars correspond to +/- standard deviation (SD).

## ACKNOWLEDGMENTS

We thank Dr. Jonathan Karn (Case Western University Reserve) for providing Jurkat E4 and 2D10 cells, and Dr. Vivian Bardwell and Dr. Micah Gearhart (University of Minnesota) for providing the ChIP-grade PCGF1 antibody. We are grateful for the technical assistance of Mélanie Laporte and Frederic Dallaire as well as for technical advice from Noe Seija Desivo from the IRCM. We thank Julie Lord from the IRCM Flow Cytometry platform. We also thank the IRCM Molecular Biology and functional genomics core laboratory staff for sequencing and assistance with cell fractionation. We are also grateful to Drs. Robert Lodge, Mariana Bego, and Javier Di Noia for technical advice. We thank Dr. Didier Trono for the WPI and PAX2 lentiviral plasmids. The following reagents were obtained through the NIH HIV Reagent Program, Division of AIDS, NIAID, NIH: Jurkat (E6-1) Cells, contributed by ATCC (Dr. Arthur Weiss), and MT4 cells, contributed by Dr. Douglas Richman. This study was supported by grants from the Canadian Institutes of Health Research (CIHR) (FDN 154324), the CIHR-supported Canadian HIV Cure Enterprise (CanCURE 2.0) (HB2 164064), and the Fonds de Recherche du Québec-Santé AIDS and Infectious Disease Network to É.A.C. M.C.R-M. is a recipient of a postdoctoral fellowship from Fonds de Recherche du Québec-Santé (#356778). É.A.C. is the recipient of the IRCM-Université de Montréal Chair of Excellence in HIV Research.

## AUTHOR CONTRIBUTIONS

Experimentation, data analysis, and writing of manuscript, M.C.R-M. and I.M-A. Generation of preliminary data and contribution to reagent development, J.A.F.B. Contribution to data analysis and manuscript review T.N.Q.P. Acquisition of funding, conception, data analysis, supervision of research and manuscript writing, E.A.C.

## DATA AVAILABILITY

All data are within the article and supplemental material

## SUPPLEMENTAL INFORMATION

Figures S1-S3.

Table 1.

## Supplemental Figure Legends

**Supplemental Figure 1**

*Vpr-mediated BCOR depletion is independent of G2/M cell cycle arrest and cytoxicity,* related to Figure 2

(A) HeLa Flp-In T-Rex cells expressing or not HA-WT-Vpr were treated with doxycycline for the indicated time, stained with Aqua Live/dead exclusion dye, and analyzed by flow cytometry. Shown is the frequency of Aqua Live dye positive cells (dead) at each time point in one of two experiments. (B) Expression of HA-WT-Vpr or the G2-arrest defective S79A-Vpr was induced in HeLa Flp-In T-Rex cells for up to 24 h. Cell cycle analysis was then performed by PI staining and flow cytometry. Shown is a representative analysis with the G2/M:G1 ratio indicated for each condition (n=2). (C) Following the induction of WT- or S79A-Vpr for the indicated time interval (left panel), protein lysates were analyzed by immunoblotting for the indicated proteins (left panel with quantification of BCOR and HLTF expression indicated below). Compiled quantitative results from two experiments are indicated (right panel).

**Supplemental Figure 2**

*Specific BCOR knock-down in Jurkat E4*, related to Figure 5

(A) Western blot of BCOR basal expression levels in 293T and 293T BCOR knockout cells as well as in Jurkat E6.1 (parental), Jurkat E4 and Jurkat 2D10 cells.

(B) Schematic representation of the HIV-1 LTR region. Numbers represent coordinates relative to the transcriptional start site (TSS= +1). Nuc 0 and Nuc 1 stand for Nucleosome 0 and Nucleosome 1 respectively. Arrows indicate the position of the primer DNA sequence encompassing, Nuc -0 (black), the transcriptional start site (purple) and Nuc-1 (orange).

(C) Schematic representation of the lentiviral vector used to express BCOR-targeting shRNAs under a U6 promoter as well as a mCherry reporter gene under the control of the Spleen Focus-Forming Virus (SFFV) promoter.

(D) A representative immunoblot of BCOR expression levels in Jurkat E4 transduced with 5 μL of the three distinct shRNAs relative to cells transduced with the scramble shRNA control. Detection of Lamin B1 (LMNB1) was used as a loading control.

**Supplemental Figure 3**

*BCOR contributes to LTR silencing in the Jurkat 2D10 model of HIV-1 latency,* related to Figure 5.

(A) Schematic representation of the integrated HIV-1 provirus in the 2D10 model cell line of HIV-1 latency. The integrated provirus expresses a Tat mutant (H13L) with an attenuated transactivating activity as well as a GFP with a short half-life (d2EGFP). (B) Histograms summarizing ChIP-qPCR results for the indicated proteins expressed as percentage of input at the HIV-1 LTR (Nuc 0, Nuc 1 or TSS) or the GAPDH promoter DNA sequences; each color represents an independent experiment (n=5).

(C) 2D10 cells were transduced with mCherry-reporter lentivirus expressing either Scramble, shBCOR#2 at the indicated input doses or the viral protein Tat at 2.5 μL. GFP expression in the transduced mCherry^+^ cells was determined by flow cytometry (left panels). A graph summarizing the analysis of three independent BCOR-depletion experiments is shown (right panel). For all panels, Error bars, SD; Statistical analysis: Ordinary one-way Anova, *p< 0.05, ***p< 0.001, ****p<0.0001.

## REFERENCES

1. Chen HC, Martinez JP, Zorita E, Meyerhans A, Filion GJ. 2017. Position effects influence HIV latency reversal. Nat Struct Mol Biol 24:47–54.

2. Jordan A, Defechereux P, Verdin E. 2001. The site of HIV-1 integration in the human genome determines basal transcriptional activity and response to Tat transactivation. EMBO J 20:1726–38.

3. Jiang C, Lian X, Gao C, Sun X, Einkauf KB, Chevalier JM, Chen SMY, Hua S, Rhee B, Chang K, Blackmer JE, Osborn M, Peluso MJ, Hoh R, Somsouk M, Milush J, Bertagnolli LN, Sweet SE, Varriale JA, Burbelo PD, Chun TW, Laird GM, Serrao E, Engelman AN, Carrington M, Siliciano RF, Siliciano JM, Deeks SG, Walker BD, Lichterfeld M, Yu XG. 2020. Distinct viral reservoirs in individuals with spontaneous control of HIV-1. Nature 585:261–267.

4. Jordan A, Bisgrove D, Verdin E. 2003. HIV reproducibly establishes a latent infection after acute infection of T cells in vitro. EMBO J 22:1868–77.

5. Tripathy MK, Abbas W, Herbein G. 2011. Epigenetic regulation of HIV-1 transcription. Epigenomics 3:487–502.

6. Siliciano JD, Kajdas J, Finzi D, Quinn TC, Chadwick K, Margolick JB, Kovacs C, Gange SJ, Siliciano RF. 2003. Long-term follow-up studies confirm the stability of the latent reservoir for HIV-1 in resting CD4+ T cells. Nat Med 9:727–8.

7. Tchasovnikarova IA, Timms RT, Matheson NJ, Wals K, Antrobus R, Gottgens B, Dougan G, Dawson MA, Lehner PJ. 2015. GENE SILENCING. Epigenetic silencing by the HUSH complex mediates position-effect variegation in human cells. Science 348:1481–1485.

8. Yurkovetskiy L, Guney MH, Kim K, Goh SL, McCauley S, Dauphin A, Diehl WE, Luban J. 2018. Primate immunodeficiency virus proteins Vpx and Vpr counteract transcriptional repression of proviruses by the HUSH complex. Nat Microbiol 3:1354–1361.

9. Chougui G, Munir-Matloob S, Matkovic R, Martin MM, Morel M, Lahouassa H, Leduc M, Ramirez BC, Etienne L, Margottin-Goguet F. 2018. HIV-2/SIV viral protein X counteracts HUSH repressor complex. Nat Microbiol 3:891–897.

10. Dupont L, Bloor S, Williamson JC, Cuesta SM, Shah R, Teixeira-Silva A, Naamati A, Greenwood EJD, Sarafianos SG, Matheson NJ, Lehner PJ. 2021. The SMC5/6 complex compacts and silences unintegrated HIV-1 DNA and is antagonized by Vpr. Cell Host Microbe 29:792–805 e6.

11. Goff SP. 2021. Silencing of Unintegrated Retroviral DNAs. Viruses 13.

12. Guenzel CA, Herate C, Benichou S. 2014. HIV-1 Vpr-a still “enigmatic multitasker“. Front Microbiol 5:127.

13. Dehart JL, Planelles V. 2008. Human immunodeficiency virus type 1 Vpr links proteasomal degradation and checkpoint activation. J Virol 82:1066–72.

14. Rogel ME, Wu LI, Emerman M. 1995. The human immunodeficiency virus type 1 vpr gene prevents cell proliferation during chronic infection. J Virol 69:882–8.

15. Belzile JP, Duisit G, Rougeau N, Mercier J, Finzi A, Cohen EA. 2007. HIV-1 Vpr-mediated G2 arrest involves the DDB1-CUL4AVPRBP E3 ubiquitin ligase. PLoS Pathog 3:e85.

16. Le Rouzic E, Belaidouni N, Estrabaud E, Morel M, Rain JC, Transy C, Margottin-Goguet F. 2007. HIV1 Vpr arrests the cell cycle by recruiting DCAF1/VprBP, a receptor of the Cul4-DDB1 ubiquitin ligase. Cell Cycle 6:182–8.

17. Ahn J, Vu T, Novince Z, Guerrero-Santoro J, Rapic-Otrin V, Gronenborn AM. 2010. HIV-1 Vpr loads uracil DNA glycosylase-2 onto DCAF1, a substrate recognition subunit of a cullin 4A-ring E3 ubiquitin ligase for proteasome-dependent degradation. J Biol Chem 285:37333–41.

18. Laguette N, Bregnard C, Hue P, Basbous J, Yatim A, Larroque M, Kirchhoff F, Constantinou A, Sobhian B, Benkirane M. 2014. Premature activation of the SLX4 complex by Vpr promotes G2/M arrest and escape from innate immune sensing. Cell 156:134–45.

19. Hrecka K, Hao C, Shun MC, Kaur S, Swanson SK, Florens L, Washburn MP, Skowronski J. 2016. HIV-1 and HIV-2 exhibit divergent interactions with HLTF and UNG2 DNA repair proteins. Proc Natl Acad Sci U S A 113:E3921–30.

20. Lahouassa H, Blondot ML, Chauveau L, Chougui G, Morel M, Leduc M, Guillonneau F, Ramirez BC, Schwartz O, Margottin-Goguet F. 2016. HIV-1 Vpr degrades the HLTF DNA translocase in T cells and macrophages. Proc Natl Acad Sci U S A 113:5311–6.

21. Yan J, Shun MC, Hao C, Zhang Y, Qian J, Hrecka K, DeLucia M, Monnie C, Ahn J, Skowronski J. 2018. HIV-1 Vpr Reprograms CLR4(DCAF1) E3 Ubiquitin Ligase to Antagonize Exonuclease 1-Mediated Restriction of HIV-1 Infection. mBio 9.

22. Greenwood EJ, Matheson NJ, Wals K, van den Boomen DJ, Antrobus R, Williamson JC, Lehner PJ. 2016. Temporal proteomic analysis of HIV infection reveals remodelling of the host phosphoproteome by lentiviral Vif variants. Elife 5.

23. Greenwood EJD, Williamson JC, Sienkiewicz A, Naamati A, Matheson NJ, Lehner PJ. 2019. Promiscuous Targeting of Cellular Proteins by Vpr Drives Systems-Level Proteomic Remodeling in HIV-1 Infection. Cell Rep 27:1579–1596 e7.

25. Forouzanfar F, Ali S, Wallet C, De Rovere M, Ducloy C, El Mekdad H, El Maassarani M, Ait-Ammar A, Van Assche J, Boutant E, Daouad F, Margottin-Goguet F, Moog C, Van Lint C, Schwartz C, Rohr O. 2019. HIV-1 Vpr mediates the depletion of the cellular repressor CTIP2 to counteract viral gene silencing. Sci Rep 9:13154.

26. Maudet C, Sourisce A, Dragin L, Lahouassa H, Rain JC, Bouaziz S, Ramirez BC, Margottin-Goguet F. 2013. HIV-1 Vpr induces the degradation of ZIP and sZIP, adaptors of the NuRD chromatin remodeling complex, by hijacking DCAF1/VprBP. PLoS One 8:e77320.

27. Lv L, Wang Q, Xu Y, Tsao LC, Nakagawa T, Guo H, Su L, Xiong Y. 2018. Vpr Targets TET2 for Degradation by CRL4(VprBP) E3 Ligase to Sustain IL-6 Expression and Enhance HIV-1 Replication. Mol Cell 70:961–970 e5.

28. Zhou X, Monnie C, DeLucia M, Ahn J. 2021. HIV-1 Vpr activates host CRL4-DCAF1 E3 ligase to degrade histone deacetylase SIRT7. Virol J 18:48.

29. Blackledge NP, Klose RJ. 2021. The molecular principles of gene regulation by Polycomb repressive complexes. Nat Rev Mol Cell Biol 22:815–833.

30. Cooper S, Grijzenhout A, Underwood E, Ancelin K, Zhang T, Nesterova TB, Anil-Kirmizitas B, Bassett A, Kooistra SM, Agger K, Helin K, Heard E, Brockdorff N. 2016. Jarid2 binds mono-ubiquitylated H2A lysine 119 to mediate crosstalk between Polycomb complexes PRC1 and PRC2. Nat Commun 7:13661.

31. Laugesen A, Hojfeldt JW, Helin K. 2019. Molecular Mechanisms Directing PRC2 Recruitment and H3K27 Methylation. Mol Cell 74:8–18.

32. Zhou W, Zhu P, Wang J, Pascual G, Ohgi KA, Lozach J, Glass CK, Rosenfeld MG. 2008. Histone H2A monoubiquitination represses transcription by inhibiting RNA polymerase II transcriptional elongation. Mol Cell 29:69–80.

33. Rose NR, King HW, Blackledge NP, Fursova NA, Ember KJ, Fischer R, Kessler BM, Klose RJ. 2016. RYBP stimulates PRC1 to shape chromatin-based communication between Polycomb repressive complexes. Elife 5.

34. Garcia E, Marcos-Gutierrez C, del Mar Lorente M, Moreno JC, Vidal M. 1999. RYBP, a new repressor protein that interacts with components of the mammalian Polycomb complex, and with the transcription factor YY1. EMBO J 18:3404–18.

35. Piunti A, Shilatifard A. 2021. The roles of Polycomb repressive complexes in mammalian development and cancer. Nat Rev Mol Cell Biol 22:326–345.

36. Friedman J, Cho WK, Chu CK, Keedy KS, Archin NM, Margolis DM, Karn J. 2011. Epigenetic silencing of HIV-1 by the histone H3 lysine 27 methyltransferase enhancer of Zeste 2. J Virol 85:9078–89.

37. Khan S, Iqbal M, Tariq M, Baig SM, Abbas W. 2018. Epigenetic regulation of HIV-1 latency: focus on polycomb group (PcG) proteins. Clin Epigenetics 10:14.

38. Nguyen K, Das B, Dobrowolski C, Karn J. 2017. Multiple Histone Lysine Methyltransferases Are Required for the Establishment and Maintenance of HIV-1 Latency. mBio 8.

39. Turner AW, Margolis DM. 2017. Chromatin Regulation and the Histone Code in HIV Latency. Yale J Biol Med 90:229–243.

40. Sharma AL, Hokello J, Sonti S, Zicari S, Sun L, Alqatawni A, Bukrinsky M, Simon G, Chauhan A, Daniel R, Tyagi M. 2020. CBF-1 Promotes the Establishment and Maintenance of HIV Latency by Recruiting Polycomb Repressive Complexes, PRC1 and PRC2, at HIV LTR. Viruses 12.

41. Tripathy MK, McManamy ME, Burch BD, Archin NM, Margolis DM. 2015. H3K27 Demethylation at the Proviral Promoter Sensitizes Latent HIV to the Effects of Vorinostat in Ex Vivo Cultures of Resting CD4+ T Cells. J Virol 89:8392–405.

42. Coull JJ, Romerio F, Sun JM, Volker JL, Galvin KM, Davie JR, Shi Y, Hansen U, Margolis DM. 2000. The human factors YY1 and LSF repress the human immunodeficiency virus type 1 long terminal repeat via recruitment of histone deacetylase 1. J Virol 74:6790–9.

43. Margolis DM, Somasundaran M, Green MR. 1994. Human transcription factor YY1 represses human immunodeficiency virus type 1 transcription and virion production. J Virol 68:905–10.

44. Gearhart MD, Corcoran CM, Wamstad JA, Bardwell VJ. 2006. Polycomb group and SCF ubiquitin ligases are found in a novel BCOR complex that is recruited to BCL6 targets. Mol Cell Biol 26:6880–9.

45. Yan J, Shun MC, Zhang Y, Hao C, Skowronski J. 2019. HIV-1 Vpr counteracts HLTF-mediated restriction of HIV-1 infection in T cells. Proc Natl Acad Sci U S A 116:9568–9577.

46. Tristem M, Marshall C, Karpas A, Hill F. 1992. Evolution of the primate lentiviruses: evidence from vpx and vpr. EMBO J 11:3405–12.

47. Planelles V, Barker E. 2010. Roles of Vpr and Vpx in modulating the virus-host cell relationship. Mol Aspects Med 31:398–406.

48. Schwefel D, Groom HC, Boucherit VC, Christodoulou E, Walker PA, Stoye JP, Bishop KN, Taylor IA. 2014. Structural basis of lentiviral subversion of a cellular protein degradation pathway. Nature 505:234–8.

49. Cohen GB, Gandhi RT, Davis DM, Mandelboim O, Chen BK, Strominger JL, Baltimore D. 1999. The selective downregulation of class I major histocompatibility complex proteins by HIV-1 protects HIV-infected cells from NK cells. Immunity 10:661–71.

50. Richard J, Sindhu S, Pham TN, Belzile JP, Cohen EA. 2010. HIV-1 Vpr up-regulates expression of ligands for the activating NKG2D receptor and promotes NK cell-mediated killing. Blood 115:1354–63.

51. Ly T, Endo A, Lamond AI. 2015. Proteomic analysis of the response to cell cycle arrests in human myeloid leukemia cells. Elife 4.

52. Zhou Y, Ratner L. 2000. Phosphorylation of human immunodeficiency virus type 1 Vpr regulates cell cycle arrest. J Virol 74:6520–7.

53. Barnitz RA, Wan F, Tripuraneni V, Bolton DL, Lenardo MJ. 2010. Protein kinase A phosphorylation activates Vpr-induced cell cycle arrest during human immunodeficiency virus type 1 infection. J Virol 84:6410–24.

54. Belzile JP, Richard J, Rougeau N, Xiao Y, Cohen EA. 2010. HIV-1 Vpr induces the K48-linked polyubiquitination and proteasomal degradation of target cellular proteins to activate ATR and promote G2 arrest. J Virol 84:3320–30.

55. Ahn J, Novince Z, Concel J, Byeon CH, Makhov AM, Byeon IJ, Zhang P, Gronenborn AM. 2011. The Cullin-RING E3 ubiquitin ligase CRL4-DCAF1 complex dimerizes via a short helical region in DCAF1. Biochemistry 50:1359–67.

56. Merlet J, Burger J, Gomes JE, Pintard L. 2009. Regulation of cullin-RING E3 ubiquitin-ligases by neddylation and dimerization. Cell Mol Life Sci 66:1924–38.

57. Huynh KD, Fischle W, Verdin E, Bardwell VJ. 2000. BCoR, a novel corepressor involved in BCL-6 repression. Genes Dev 14:1810–23.

58. Pearson R, Kim YK, Hokello J, Lassen K, Friedman J, Tyagi M, Karn J. 2008. Epigenetic silencing of human immunodeficiency virus (HIV) transcription by formation of restrictive chromatin structures at the viral long terminal repeat drives the progressive entry of HIV into latency. J Virol 82:12291–303.

59. Chougui G, Margottin-Goguet F. 2019. HUSH, a Link Between Intrinsic Immunity and HIV Latency. Front Microbiol 10:224.

60. Mbonye U, Karn J. 2024. The cell biology of HIV-1 latency and rebound. Retrovirology 21:6.

61. Cohen EA, Dehni G, Sodroski JG, Haseltine WA. 1990. Human immunodeficiency virus vpr product is a virion-associated regulatory protein. J Virol 64:3097–9.

62. Planelles V, Jowett JB, Li QX, Xie Y, Hahn B, Chen IS. 1996. Vpr-induced cell cycle arrest is conserved among primate lentiviruses. J Virol 70:2516–24.

63. Xu W, Wu Y, Zhao J, Chen J, Zhang W. 2020. Human immunodeficiency virus type 1 transcription is regulated by thieno[3,4-d]pyrimidine. Exp Ther Med 19:3090–3096.

64. Wang J, Yang J, Yang Z, Lu X, Jin C, Cheng L, Wu N. 2016. RbAp48, a novel inhibitory factor that regulates the transcription of human immunodeficiency virus type 1. Int J Mol Med 38:267–74.

65. du Chene I, Basyuk E, Lin YL, Triboulet R, Knezevich A, Chable-Bessia C, Mettling C, Baillat V, Reynes J, Corbeau P, Bertrand E, Marcello A, Emiliani S, Kiernan R, Benkirane M. 2007. Suv39H1 and HP1gamma are responsible for chromatin-mediated HIV-1 transcriptional silencing and post-integration latency. EMBO J 26:424–35.

66. Dewhurst S, Embretson JE, Anderson DC, Mullins JI, Fultz PN. 1990. Sequence analysis and acute pathogenicity of molecularly cloned SIVSMM-PBj14. Nature 345:636–40.

67. Hsu PD, Scott DA, Weinstein JA, Ran FA, Konermann S, Agarwala V, Li Y, Fine EJ, Wu X, Shalem O, Cradick TJ, Marraffini LA, Bao G, Zhang F. 2013. DNA targeting specificity of RNA-guided Cas9 nucleases. Nat Biotechnol 31:827–32.

68. Pham TN, Richard J, Gerard FC, Power C, Cohen EA. 2011. Modulation of NKG2D-mediated cytotoxic functions of natural killer cells by viral protein R from HIV-1 primary isolates. J Virol 85:12254–61.

69. Hrimech M, Yao XJ, Bachand F, Rougeau N, Cohen EA. 1999. Human immunodeficiency virus type 1 (HIV-1) Vpr functions as an immediate-early protein during HIV-1 infection. J Virol 73:4101–9.

